# Theta phase synchronization between the human hippocampus and the prefrontal cortex supports learning of unexpected information

**DOI:** 10.1101/144634

**Authors:** Matthias J. Gruber, Liang-Tien Hsieh, Bernhard P. Staresina, Christian E. Elger, Juergen Fell, Nikolai Axmacher, Charan Ranganath

## Abstract

Events that violate predictions are thought to not only modulate activity within the hippocampus and prefrontal cortex, but also to enhance communication between the two regions. Several studies in rodents have shown that synchronized theta oscillations facilitate communication between the prefrontal cortex and hippocampus during salient events, but it remains unclear whether similar oscillatory mechanisms support interactions between the two regions in humans. Here, we had the rare opportunity to conduct simultaneous electrophysiological recordings from the human hippocampus and prefrontal cortex from two patients undergoing presurgical evaluation for pharmaco-resistant epilepsy. Recordings were conducted during a task that involved encoding of contextually expected and unexpected visual stimuli. Across both patients, hippocampal-prefrontal theta phase synchronization was significantly higher during encoding of unexpected study items, compared to contextually expected study items. In contrast, we did not find increased theta synchronization between the prefrontal cortex and rhinal cortex. Our findings are consistent with the idea that theta oscillations orchestrate communication between the hippocampus and prefrontal cortex during the processing of contextually salient information.

## Main text

Unexpected events that violate internal predictions are more likely to be successfully encoded to memory. It has been proposed that the hippocampus and the prefrontal cortex (PFC) play a critical role in the detection and formation of memories of unexpected events (e.g., Ranganath & Rainer, 2003; Lisman & Grace, 2005). Consistent with this idea, local field potential recordings in rodents have shown that salient events (e.g. those occurring at choice points in a maze learning task) increase oscillatory power in the theta band (4-8 Hz) within the hippocampus and the PFC (e.g., Winson, 1978; Hasselmo et al., 2002; O’Neill et al., 2013; Totah et al., 2013; Donnelly et al., 2014). In addition, studies in humans have shown increases in theta oscillations at frontal electrode sites via scalp-recorded electroencephalography (EEG) and in the hippocampus via intracranial EEG recordings (e.g., Ekstrom et al., 2005; Axmacher et al., 2010; Chen et al., 2013; Gruber et al., 2013; Hsieh & Ranganath, 2014; Long et al., 2014).

Consistent with the idea that theta oscillations facilitate communication between the PFC and hippocampus, recordings in rodents and non-human primates have also shown synchronized theta oscillations between the two areas (Benchenane et al., 2010; Brincat & Miller, 2015; Fujisawa & Buzsáki, 2011; Hyman et al., 2005; Jones & Wilson, 2005). For example, enhanced theta phase synchrony between the hippocampus and the PFC has been shown during performance of a spatial T-maze task (Benchenane et al., 2010) and during retrieval of object-context associations (Place et al., 2016). These studies provide evidence that, in the rodent brain, interactions between the hippocampus and PFC rely on theta synchrony.

Little is known about the extent to which the findings of frontal-hippocampal synchronization in rodents correspond to activity in the human brain. Some intracranial EEG studies have reported increased theta phase synchronization between the PFC and neocortical regions in the medial temporal lobes (Kahana et al., 1999; Anderson et al., 2010; Watrous et al., 2013; in contrast, see Raghavachari et al., 2006), but these studies did not report changes in phase synchrony specifically with the hippocampus.

In the present study, we used intracranial EEG to determine whether human hippocampal-PFC theta phase synchrony is enhanced during processing of contextually salient, unexpected events. We recorded intracranial EEG simultaneously from the hippocampus and PFC in two pharmaco-resistant epilepsy patients while they encoded contextually expected and unexpected items. The locations of the implanted prefrontal electrodes also allowed us to explore whether theta phase synchronization with the hippocampus might be evident with specific subregions of the PFC. In addition, we also investigated phase synchronization between the PFC and sites in the rhinal cortex.

We recorded intracranial EEG from two pharmaco-refractory epileptic patients at the Department of Epileptology at the University of Bonn, Germany. Both patients (one female; 46 and 48 of age) were implanted with bilateral depth electrodes in the hippocampus and its surrounding medial temporal lobe (MTL), as well as with bilateral subdural electrodes covering parts of the PFC (i.e. one fronto-polar and one fronto-lateral electrode strip bilaterally covering rostral/ anterior and lateral PFC regions, respectively; see Fig. 1). Details about the patients and analyses of event-related potentials and oscillatory power from hippocampal sites in these two patients are presented in Axmacher et al. (2010). Because epileptic seizures were focused on left hippocampal and temporo-mesial areas in one patient and left temporo-mesial and left temporo-lateral areas in the other patient, we only considered data from the hippocampal, rhinal and PFC electrodes on the right hemisphere. The local ethics committee approved the study, and both patients gave written informed consent.

**Figure 1.**
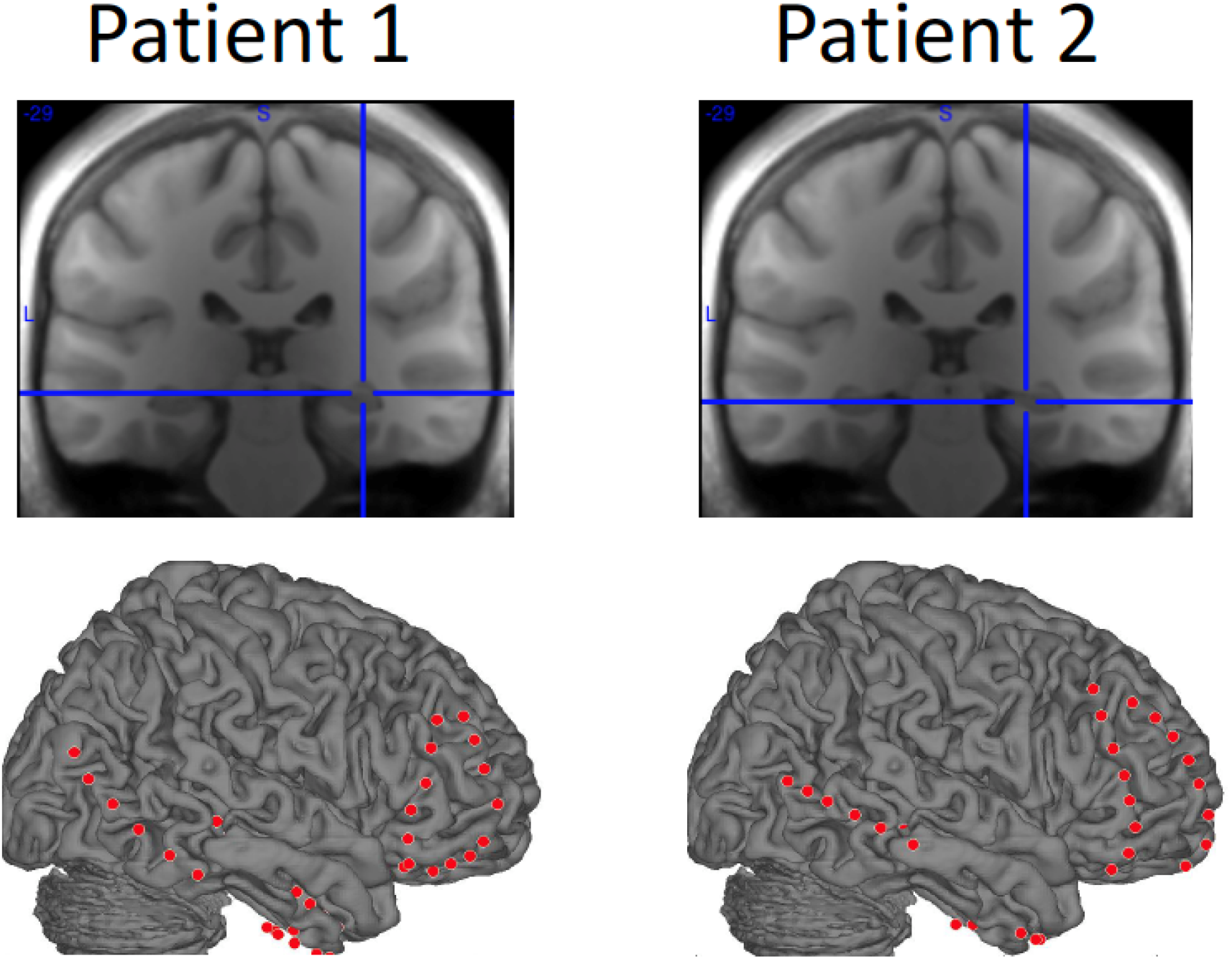
Locations of hippocampal electrode and prefrontal electrodes. On the top, the location of the selected hippocampal electrode is depicted for each patient (Patient 1: MNI 32 −29 −7; Patient 2: MNI 26 −29 −10). On the bottom, all implanted subdural strip electrodes covering the right hemisphere are depicted for each patient. Only the frontopolar and frontolateral strips were analyzed for each patient.

Both patients took part in a variant of a “Von Restorff” paradigm (Von Restorff, 1933; for details of the experimental procedure, see Axmacher et al., 2010). During the encoding phase for which iEEG results are reported here, patients encoded trial-unique images from two different categories (for exemplary trials, see Fig. 2a). Importantly, one type of stimuli comprised of the majority of encoding stimuli in a given encoding-test block (i.e. “contextually expected items”; e.g. grayscale faces on a red background as shown in Fig. 2a), and the other type of stimuli only comprised a small percentage (i.e. 14%) of the encoding stimuli in a given encoding-test block (i.e. “contextually unexpected items”; e.g. grayscale houses on a green background as shown in Fig. 2a). Categories and colors of expected and unexpected stimuli were counterbalanced across blocks in each patient. Following the encoding phase, patients completed a recognition memory test for these images (Fig. 2b).

**Figure 2.**
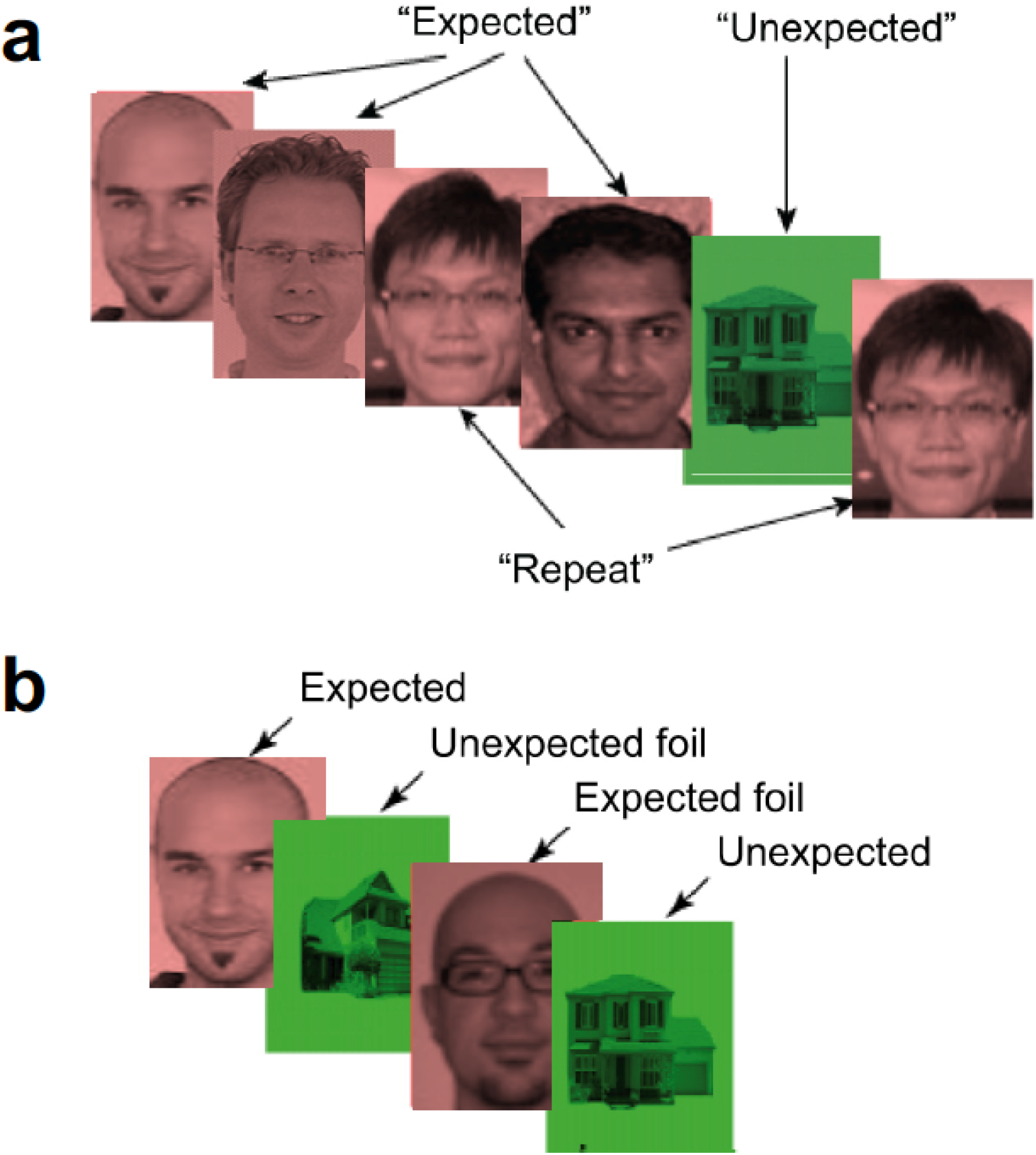
Experimental Procedure. (a) During the encoding phase for which iEEG results are reported here, patients encoded images of stimuli that comprised of the majority of encoding stimuli (“expected items”) and the other type of stimuli only comprised a small percentage (“unexpected items”). (b) Following an encoding block, patients completed a recognition memory test.

Because we were interested in how theta oscillations modulate the encoding of expected and unexpected stimuli, we restricted all iEEG analyses to ‘unexpected’ (Patient 1: 32 trials; Patient 2: 15 trials) and ‘expected’ (Patient 1: 68 trials; Patient 2: 45 trials) items that were later correctly recognized in the recognition memory test (i.e. collapsed across correct ‘confident old’ and ‘unconfident old’ responses). Because electrode placement varied across patients due to clinical needs of each patient, we focused our analyses on hippocampal contacts that were most consistently localized across the two patients. That is, we first selected one hippocampal electrode per patient that had maximal anatomical overlap between the two patients. The selected hippocampal electrode pair (one electrode from each patient) had the smallest Euclidian distance between the two patients (7 mm distance; Patient 1: MNI 32 −29 −7; Patient 2: MNI 26 −29 −10; see Fig. 1). We then visually inspected the hippocampal and prefrontal raw data and excluded all data of the first electrode on each right PFC electrode strip (i.e. most-inferior electrode) due to a very low signal-to-noise ratio as compared to all other remaining PFC electrodes. We then used the EEGLAB toolbox (Delorme & Makeig, 2004) to epoch the iEEG data into segments from −2s to +3s relative to the presentation onset of all items. Trials containing artifacts were then manually discarded from the analyses. Artifact-free iEEG data were then imported into the Fieldtrip toolbox (Oostenveld et al., 2011) for our analyses of interests. Within the Fieldtrip toolbox, first the artifact-free raw EEG data underwent standard time-frequency decomposition analysis to obtain power and phase information (i.e. Morlet wavelet with a width of 5 cycles; time period: −0.5s to 1.5s in steps of 0.02s; frequency band: 2-30 Hz). Second, in order to address the role of theta phase synchrony between the hippocampus and the PFC, we calculated phase synchrony indices between the previously selected hippocampal electrode and each of the artifact-free frontal electrodes, resulting in 14 hippocampal-PFC electrode pairs for each patient. Phase synchrony was separately quantified for unexpected and expected trials using the weighted phase lag index (WPLI) implemented in Fieldtrip. The WPLI has the advantage that it alleviates problems related to volume conduction and other noise-related issues (Vinck et al., 2011). For statistical analyses, we calculated an averaged WPLI value based on an a-priori selected theta time-frequency bin for each hippocampal-PFC electrode pair (i.e. average of 4-5 Hz in 200-400ms after stimulus onset). The 4-5 Hz frequency bin was selected based on the maximum difference in spectral power between unexpected and expected items in the selected hippocampal electrodes. The time window was selected based on our previous results (Axmacher et al., 2010), which revealed significant increases in theta power 200-400 ms following encoding of unexpected as compared to expected items. Interestingly, visual inspection of all electrode pairs during this time-frequency bin suggested that several hippocampal-PFC electrode pairs showed increased theta phase synchrony for unexpected, but not for expected items (see Fig. 3 for selected electrodes showing significant findings that survived multiple comparisons correction).

**Figure 3.**
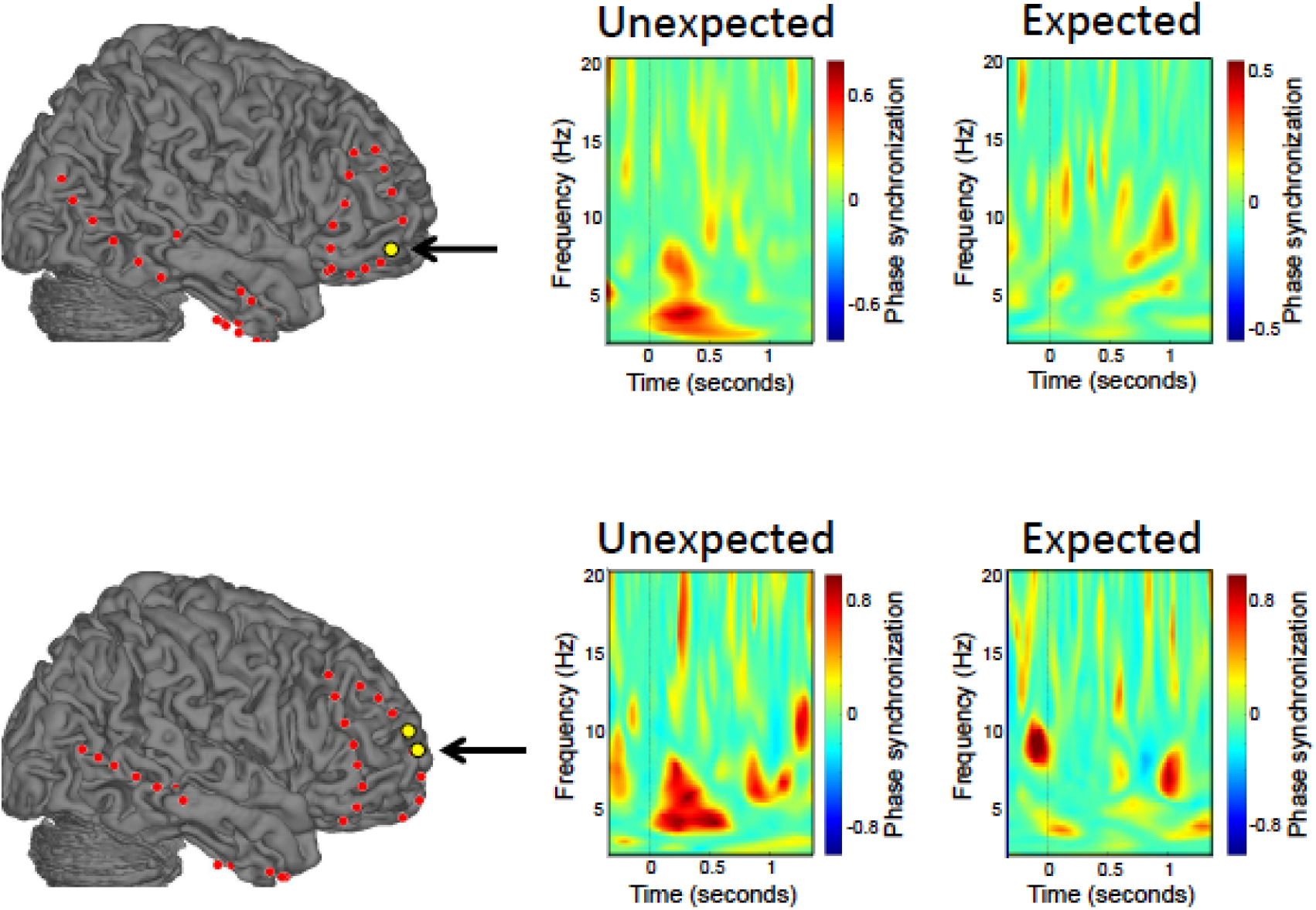
Increases in hippocampal-prefrontal theta phase synchrony for unexpected compared to expected information. In both patients, frontopolar electrode sites showed significantly increased theta phase synchrony with the hippocampus during encoding of unexpected compared to expected items. Electrodes with between-condition differences that survived multiple-comparisons correction are highlighted in yellow. Arrows indicate the electrodes for which we show time-frequency spectrograms individually for unexpected and expected items. Top row depicts findings for Patient 1 and bottom row depicts findings for Patient 2.

To statistically determine whether unexpected compared to expected items show a significant theta phase synchrony increase in our selected time-frequency bin, we used a non-parametric statistical approach that randomly permutes condition labels to correct for multiple comparisons on a cluster level (i.e. across electrodes). Analyses were conducted separately in each patient. The steps are as follows: (1) We computed the averaged WPLI value in our time-frequency bin for each condition (in order to use an identical approach as for the surrogate data, we selected equal trial numbers from both conditions based on the minimum number of trials in one condition). We then computed the difference of the averaged WPLI value between the unexpected and the expected items in order to obtain the empirical difference in theta phase synchrony (i.e. WPLI) between conditions. (2) We shuffled trial labels by randomly selecting equal trial numbers from both conditions based on the minimum number of trials in one condition, calculated surrogate phase synchrony values for all 14 electrode pairs, took the difference between the surrogate conditions for all 14 electrode pairs, and saved the maximum surrogate phase synchrony difference across all 14 electrode pairs (i.e. electrode-pair_max_). (3) Step 2 was repeated 500 times. Based on the 500 permutations, we created a null distribution of all electrode-pair_max_ difference values and determined the alpha cut-off point (*p* < 0.05; one-sided; i.e. 475^th^ data point in surrogate difference distribution) in order to test the statistical significance of the empirical theta phase synchrony values for all electrode pairs. This stringent approach allowed us to correct for multiple comparisons across electrodes.

As shown in Fig. 3, in both patients, frontopolar electrode sites showed significantly increased theta phase synchrony with the hippocampus during encoding of unexpected compared to expected items. To examine the spatial specificity of the observed hippocampal-PFC theta phase synchrony effect, we performed control analyses in which we quantified theta phase synchrony between rhinal and PFC electrodes. Consistent with the selection of the hippocampal electrodes, we also selected a spatially close electrode contact for each patient from the rhinal cortex (perirhinal/ entorhinal cortex) based on the smallest Euclidian distance between rhinal contacts in both patients resulting in 9 mm distance between both patients (distance between rhinal and hippocampal contact: 41 and 36 mm for Patient 1 and 2, respectively). Importantly, even without correcting for multiple comparisons, we did not find evidence of enhanced theta phase synchrony between any rhinal-PFC electrode pairs for unexpected compared to expected trials.

In summary, results from our study demonstrate that theta phase synchrony between the hippocampus and PFC is enhanced during unexpected, contextually deviant events. Moreover, results from both participants converged in revealing that synchronization with hippocampal theta occurred at sites in the frontopolar cortex. These findings are consistent with the idea that theta oscillations facilitate communication between the PFC and hippocampus.

Although initial electrophysiological recording studies in rodents and non-human primates have provided evidence for theta synchronization between the hippocampus and PFC for salient events (Benchenane et al., 2010; Brincat & Miller, 2015; Fujisawa & Buzsáki, 2011), it is worth noting that non-human and human electrophysiological studies typically assess synchrony in different ways. Studies in rodents often measure synchrony via single-unit spiking activity that is phase-locked to theta oscillations or via amplitude-based coherence of local field potentials between two regions (e.g., Jones & Wilson, 2005; Benchenane et al., 2010), whereas human studies commonly measure synchrony via phase alignment of theta oscillations between distant brain regions (e.g., Backus et al., 2016; Kaplan et al., 2017; Watrous et al., 2013). Despite these methodological differences in the measurement of synchrony, our findings in humans converge with findings in rodents in that they support the idea that theta synchrony facilitates interactions between the hippocampus and PFC.

It could be argued that theta synchronization might be a ubiquitous phenomenon during encoding, but at least two aspects of our findings are not consistent with this idea. First, theta synchrony between the two regions was larger for unexpected compared to expected events and, second, this synchrony increase was specific between the PFC and the hippocampus, but did not extend to a cortical MTL region (i.e. no evidence for rhinal-PFC theta synchrony). Therefore, our findings suggest that increased theta synchrony might be specific to a brain network (involving the PFC and hippocampus) that detects the salience of information rather than being a ubiquitous property during encoding.

The increase in theta phase synchrony was transient and emerged during an early time period during the presentation of an unexpected event. This early theta hippocampal-PFC synchrony coincides with our previously shown early event-related potential (ERP) finding in the human hippocampus (Axmacher et al., 2010). Therefore, the increase in theta synchrony between the PFC and hippocampus together with this early hippocampal ERP might suggest an early detection process that is elicited when expectations are violated and on-going encoding processes need to be flexibly adopted towards the unexpected information (cf. Axmacher et al., 2010).

One limitation of the current study is that, in these patients, we did not have sufficient numbers of subsequently forgotten trials to adequately compare phase synchronization between subsequently remembered and subsequently forgotten items. Furthermore, only two of our patients had electrodes placed in both the hippocampus and PFC. It would be beneficial for future studies to investigate this question with a larger sample and sufficient numbers of trials to test for differences between subsequently remembered and subsequently forgotten items.

In conclusion, we have shown that contextually salient information elicits increased theta phase synchrony between the hippocampus and frontopolar cortex. Consistent with the literature on the relationship between theta activity and memory (e.g., Jones & Wilson, 2005; Watrous et al., 2013; for a review, see Hsieh & Ranganath, 2014), we suggest that theta synchrony between the hippocampus and the PFC may be a neural mechanism that helps to prioritize encoding of novel, salient information.

